# Inhibition Resolves Simon Conflict: Evidence from EEG Decoding

**DOI:** 10.1101/2024.10.13.618127

**Authors:** Yoon Seo Lee, Gi-Yeul Bae, Yang Seok Cho

## Abstract

The congruency sequence effect (CSE), a hypothesized marker of top-down cognitive control, refers to a reduced congruency effect after incongruent trials compared to congruent trials. Although this effect has been observed across various distractor interference tasks, the nature of the control processes underlying the CSE remains a topic of active debate. It has been suggested that cognitive control may resolve conflicts in information processing either by (a) enhancing the representation of goal information and/or (b) suppressing the representation of distractor information. The present study aimed to identify the conflict resolution processes within the context of the color Simon task by decoding the goal and distracting information from human scalp EEG signals. For the decoding analysis, models were trained separately for color and location attributes corresponding to goal and distractor information. Additionally, decoding accuracy was calculated in different frequency bands: theta (4-8 Hz), alpha (8-12 Hz), low beta (12-20 Hz), and high beta (20-30 Hz). Results showed that decoding accuracy for distractor information was reduced when cognitive control was activated and this pattern was only observed in the high beta-frequency band (20-30 Hz). In contrast, no such difference was observed for target information. These findings suggest that cognitive control regulates Simon conflict by inhibiting distractor representation in the brain, thereby preventing unwanted distraction-driven behaviors.

## Introduction

Cognitive control refers to the ability to regulate one’s thoughts and behaviors to adapt to external environmental demands or internal goals. It encompasses three key components: working memory, which enables information retention; set-shifting, which allows flexible attention shifts; and inhibition (or facilitation), which helps suppress automatic or dominant responses (Miyake et al., 2000). Among these components, inhibition is particularly crucial for self-regulation, as it allows individuals to override impulsive or habitual responses and stay focused on their goals (Diamond, 2013). Without inhibition, individuals are more likely to be driven by impulses, established habits, and environmental stimuli, leading to behavioral challenges and learning disabilities (Munakata et al., 2011). A comprehensive understanding of the mechanisms underlying inhibition is thus critical not only for improving clinical interventions but also for advancing insights into complex intelligent behavior.

One way to examine inhibition is using distractor interference tasks, where participants are required to make a novel response to a goal (task-relevant features) in the presence of a distractor (task-irrelevant features) that is strongly associated with a habitual response (Miyake et al., 2000). For example, in the Simon task, where participants are to respond to a target’s color, presented either to the left or right of a fixation cross, while ignoring its location. Typically, performance becomes slower and less accurate when the stimulus position and the correct response location are incongruent than when they are congruent, a phenomenon known as the congruency effect. Importantly, this congruency effect is reduced after experiencing incongruent trials compared to congruent trials (Gratton et al., 1992) – a phenomenon called the congruency sequence effect (CSE; Figure 1). The robustness of the CSE has been demonstrated across various distractor interference tasks, including the Simon task (Lee & Cho, 2013; Lim & Cho, 2021b; Notebaert & Verguts, 2008; Stürmer et al., 2002), the Stroop task (Egner & Hirsch, 2005; Kerns et al., 2004; Notebaert et al., 2006), the flanker compatibility task (Kim & Cho, 2014; Lim & Cho, 2018), and the prime-probe task (Grant et al., 2020; Grant et al., 2023).

**Figure 1.**
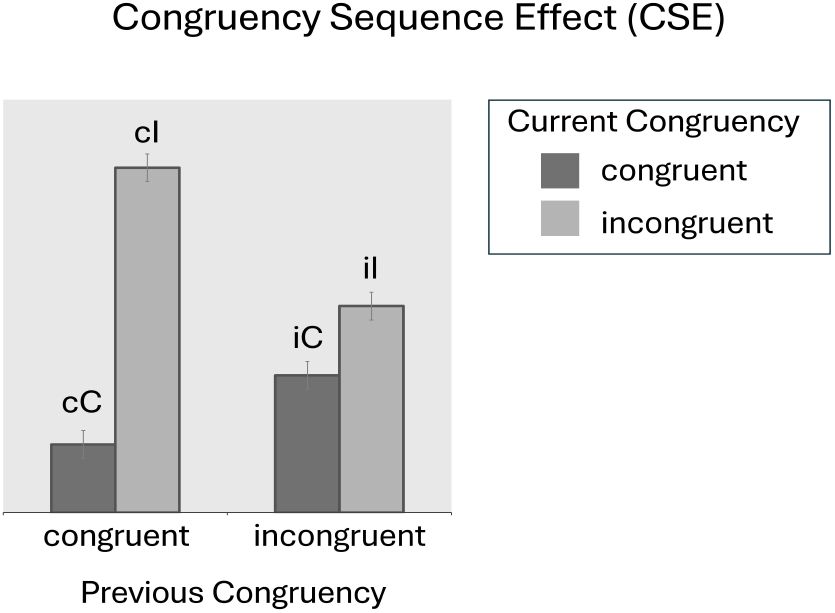
The diagram illustrates the congruency sequence effect (CSE). The first two bars represent trials where the previous congruency was congruent, while the second two bars represent trials where the previous congruency was incongruent. In each set, the dark bars correspond to current congruent trials, and the light bars correspond to current incongruent trials. This creates four conditions: cC (congruent after congruent), cI (incongruent after congruent), iC (congruent after incongruent), and iI (incongruent after incongruent). The CSE is characterized by current trial interference reduced after experiencing interference trials. For example, responses are typically slower for incongruent trials compared to congruent trials. However, this effect is modulated by the congruency of the previous trial. Specifically, the distraction caused by current incongruent trials is diminished after experiencing incongruent trials compared to congruent ones (i.e., iI vs. cI), reflecting the effects of conflict resolution.

According to the conflict monitoring theory (Botvinick et al., 2001), the CSE reflects increased top-down cognitive control. When response conflict occurs during an incongruent trial, the anterior cingulate cortex detects the conflict and sends signals to the dorsolateral prefrontal cortex to enhance cognitive control, thereby reducing response conflict in subsequent trials. While the top-down cognitive control accounts effectively explain the CSE, alternative views suggest that the CSE may stem from bottom-up factors, such as stimulus and response feature-repetition priming (Hommel et al., 2004; Mayr et al., 2003) or contingency learning between task-irrelevant stimulus feature and their frequently associated responses (Schmidt, & De Houwer, 2011). Nonetheless, a number of studies have consistently reported CSEs, even when both repetition-priming and contingency-learning confounds were controlled (Kim & Cho, 2014; Lim & Cho, 2021a; Schmidt & Weissman, 2014), indicating that top-down cognitive control significantly contributes to the observed CSE.

An important question regarding top-down cognitive control in the CSE is how exactly the cognitive control in the brain operates to resolve conflicts. According to the dual-route model, information processing in the Simon task occurs along two different routes (de Jong et al., 1994; Kornblum et al., 1990). The task-relevant feature is processed along a controlled route that intentionally links the stimulus to the required response, whereas the task-irrelevant feature is processed along an automatic route that directly activates the corresponding spatial responses. The overall decision is then based on a merger of both information sources (Pashler, 1997). Thus, response conflict arises when task-relevant and task-irrelevant information activate different responses. To resolve the conflict, cognitive control may operate in two possible ways: (a) by facilitating information processing through the controlled route, and/or (b) by inhibiting information processing through the automatic route (Koob et al., 2023).

Notebaert and Verguts (2008) support the facilitation account, demonstrating that the scope of the Simon-type CSE is determined by the task-relevant dimension. They found that the CSE transferred across two Simon-type tasks only when both tasks shared the same task-relevant dimension (i.e., when participants responded to orientation in both tasks), and not when the tasks involved different task-relevant dimensions (i.e., responding to color in one task and orientation in the other). Thus, the authors suggested that after an incongruent trial, cognitive control enhances attention to the perceptual features of the target, thereby reducing the impact of distractions (Egner & Hirsch, 2005; Notebaert & Verguts, 2008). Conversely, others have supported the inhibition account (Y. Lee & Cho, 2023; Stürmer et al., 2002). Stürmer et al. analyzed lateralized readiness potentials in the Simon task and observed early location-based priming effects over the motor cortex only after congruent trials, but not after incongruent trials. They suggested that after incongruent trials, response suppression is engaged to prevent the transmission of spatial codes from the automatic route to the response output, thereby reducing the influence of task-irrelevant stimulus information. While these studies provide important insights into the nature of cognitive control processes, response times and averaged evoked potentials mainly reflect the consequence of control processes rather than directly revealing how the brain modulates task-relevant and task-irrelevant information.

One possible way to address this issue is to use methods that directly measure the brain representation of task-relevant and task-irrelevant information. Electroencephalogram (EEG) based decoding is a novel statistical approach that used machine learning to analyze signal distributions across the scalp, providing insights into how the strength of information representation in the brain dynamically changes over time during task performance (Fahrenfort et al., 2018). Research suggested that various stimulus attributes can be decoded, including orientation (Bae & Luck, 2018), spatial location (Fahrenfort et al., 2018), color (Bae & Chen, 2024), and facial features (Bae, 2021). Moreover, recent research showed that the decoding of stimulus attributes in the current trial can vary based on previous trials (Bae & Luck, 2019; Bae, 2021). For instance, Bae and Luck (2019) demonstrated that information from preceding trials can be reactivated. They found that the orientation of a prior stimulus was decoded on the current trials. Similarly, Bae (2021) found that information from a previous trial influenced the perception of face identity and expression in the current trial, with decoding accuracy for the prior face identity increasing shortly after the current stimulus onset. These findings suggest that EEG-based decoding captures not only stimulus attributes on the current trial but also the stimulus attributes that were influenced by previous experience.

In the present study, we aimed to decode the human scalp EEG recordings of both task-relevant and task-irrelevant stimulus information following congruent and incongruent trials to examine the nature of cognitive control process. Specifically, we examined whether conflict resolution in the Simon task arises from enhanced processing of task-relevant stimulus information or from the suppression of task-irrelevant stimulus information. Participants were instructed to perform horizontal and vertical color Simon tasks alternatively in a trial-by-trial manner (Figure 2A). Decoding analysis involved training separate models to decode color and location information for task-relevant and task-irrelevant stimulus attributes, respectively. We hypothesized that if conflict resolution enhances the processing of goal-related information, decoding accuracy for task-relevant stimulus attributes will improve following incongruent trials compared to congruent trials. Conversely, if conflict resolution suppresses distractor processing, decoding accuracy for task-irrelevant stimulus attributes will decrease following incongruent trials compared to congruent trials.

**Figure 2.**
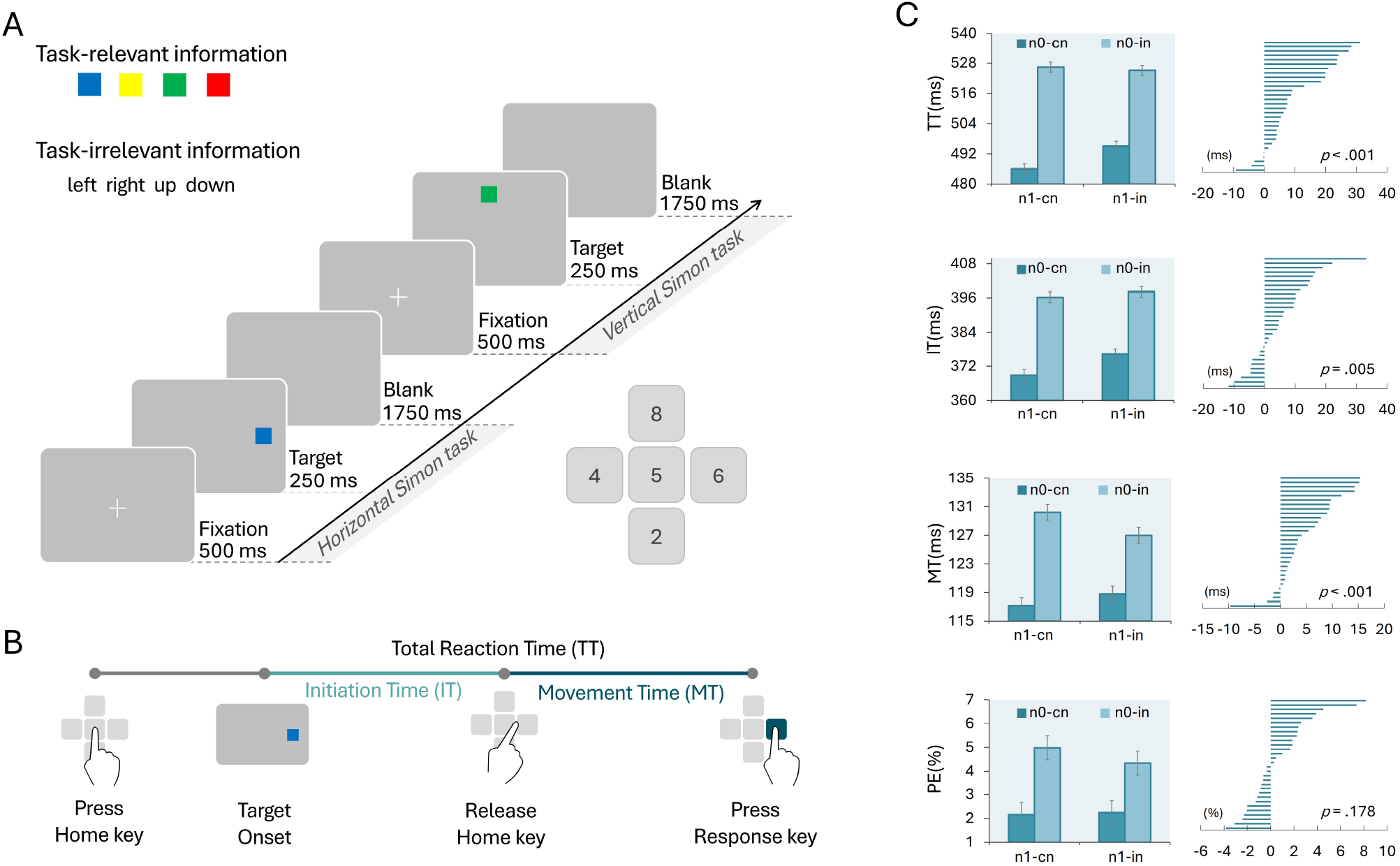
Task paradigm and behavioral results. A. An example of task sequence and response keys. Horizontal and vertical Simon tasks were presented alternatively in a trial-by-trial manner. Participants were asked to respond to the color of the square while ignoring the location of the square. Thus, the task-relevant information was color (blue, yellow, green, and red), and the task-irrelevant information was location (left, right, up, and down). B. A diagram for response method. Participants were instructed to keep pressing the home key and release it when they decided their response. Once they release the home key, they were asked to press a response key. Total reaction time (TT) was separated into initiation time (IT) and movement time (MT). IT is the temporal duration from the target onset to the release of the home key. MT is the duration from the release of the home key to the pressing of an actual response key. C. Behavioral results of TT, IT, MT, and Percentage errors (PE). Bar graphs to the left display averaged data for the four conditions cC, cI, iC, and iI. “n1-cn” refers to the previous congruent trial, “n1-in” refers to the previous incongruent trial, “n0-cn” refers to the current congruent trial, and “n0-in” refers to the current incongruent trial. Right, individual CSE = (cI - cC) - (iC - iI) were displayed.

Moreover, as previous studies have shown that decoding in different frequency bands is linked to distinct cognitive states (de Vries et al., 2019; de Vries et al., 2021; Li et al., 2023; van Driel et al., 2019), the present study calculated decoding accuracy separately for each frequency band. Numerous studies using the Simon task suggest that cognitive control is linked to theta oscillations (Cao et al., 2017; Cohen & Donner, 2013; Cohen & Ridderinkhof, 2013; Gulbinaite et al., 2014; Töllner et al., 2017; van Driel et al., 2015; Wang et al., 2019), while other have identified associations with alpha (Arnau et al., 2021; Chinn et al., 2018) or beta oscillations (Beatty et al., 2021; Duprez et al., 2019; van Es et al., 2020; Wiesman et al., 2020). However, many of these studies utilized only two response alternatives (e.g., left and right keys), which could confound the results with bottom-up repetition priming. Although some studies attempted to control for these factors by excluding stimulus repeating trials from analysis, it remains unclear whether consistent results would be obtained using a paradigm specifically designed to minimize such confounds.

To investigate top-down cognitive control, the present study adopted a bottom-up confound-minimized design. Decoding accuracy was measured in theta (4-8 Hz), alpha (8-12 Hz), low beta (12-20 Hz), and high beta (20-30 Hz) frequency bands to confirm findings from this confound-minimized design. The gamma frequency band (30-100 Hz), which is primarily associated with bottom-up processing (Buschman & Miller, 2007), was excluded from the analysis, as the study focus was solely on top-down cognitive control. Finally, a general analysis of oscillatory activity was conducted to validate the decoding analysis. We hypothesized that if conflict is resolved by enhancing attention towards task-relevant color information (Notebaert & Verguts, 2008), modulation would occur in the alpha frequency band, which has been strongly associated with attentional modulation (Foster & Awh, 2019; Peylo et al., 2021; Schneider et al., 2020; Thut et al., 2006; Van Diepen et al., 2019). If Simon-type conflict is resolved by suppressing task-irrelevant location information, specifically the automatic route from task-irrelevant dimensions to response (Stürmer et al., 2002), modulation should appear in the beta frequency band, as beta oscillations are closely linked to response modulation (Engel & Fries, 2010; Pavlidou et al., 2014; Schmidt et al., 2019; Spitzer & Haegens, 2017). Lastly, if modulation occurs in the theta frequency band, it may imply that conflict is resolved beyond simple inhibition or facilitation, as theta oscillations have been associated with working memory (Riddle et al., 2020), episodic memory (Nyhus & Curran., 2010), and episodic retrieval (Richard et al., 2004).

## Method

### Participants

The sample size was determined using G*Power 3.1 (Faul et al., 2009). Based on the experiment conducted by Y. Lee and Cho (2023, Experiment 2), where 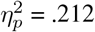, a repeated-measures analysis of variance (ANOVA) was conducted to examine the two-way interaction between previous-trial congruency (congruent or incongruent) and current-trial congruency (congruent or incongruent). The statistical power (1-*β*) set at .95, and the alpha level at 5% were applied. Our estimation indicated that a minimum sample size of 26 would provide 96% power to observe a CSE between two tasks.

Thirty participants (14 females, 16 males; mean age 23.8 years) from Korea University participated in the experiment. All participants self-reported as right-handed and no deficit in visual acuity or color vision. Prior to the experiment, all participants provided informed consent and received compensation of KRW 50,000 (about thirty-seven US dollars) after their participation. The experiment was approved by the institutional review board of Korea University (KU-IRB-16-142-A-1).

### Stimuli and apparatus

The experiment was conducted using MATLAB software (version 2015a) and the Psychophysics Toolbox version 3 (PTB-3). Visual stimuli were presented on a 24-inch (16:9) LCD monitor positioned approximately 55 cm from the participants. The sequence of displays and time course for the Simon tasks are illustrated in Figure 2A. A white (R = 255, G = 255, B = 255) cross (0.3° x 0.3° of visual angle) served as the fixation point, appearing at the center of the display. The experiment consists of vertical and horizontal Simon tasks. For the horizontal Simon task, a blue (R = 0, G = 0, B = 255) or yellow (R = 255, G = 255, B = 0) square (approximately 1.51° x 1.51°) was presented to either the left side or right side of the fixation cross. For the vertical Simon task, a red (R = 255, G = 0, B = 0) or green (R = 0, G = 255, B = 0) square was presented either above or below the fixation cross. The target stimulus appeared at an equal distance from the center of the display (approximately 5.4°). All stimuli were displayed on the gray background (R = 128, G = 128, B = 128). The aimed movement responses (Lim & Cho, 2021b) were recorded with a standard 101-key computer keyboard. The “4”, “6”, “8”, and “2” keys on the numeric keypad were used as directional responses, indicating ‘left’, ‘right’, ‘up’, and ‘down’, respectively. The “5” key or home key, was used as the starting position.

### Task design

A confound minimized design was employed (Kim & Cho, 2014; Lim & Cho, 2021a; Schmidt & Weissman, 2014). To avoid bottom-up feature integration, horizontal and vertical color Simon tasks were alternated in a trial-by-trial manner (Figure 2A). Horizontal stimulus (i.e., blue and yellow) and response sets (i.e., left and right) were presented in odd trials, while vertical stimulus (i.e., red and green) and response sets (i.e., up and down) in even trials, thereby preventing stimulus and response repetition in consecutive trials. To avoid contingency learning confounds, 8 unique stimuli (i.e., blue-left, blue-right, yellow-left, yellow-down, red-up, red-down, green-up, green-down) were presented exactly 10 times in each block.

Counterbalancing was implemented to ensure that decoding focused exclusively on task-relevant color attributes, without being influenced by specific response associations. Furthermore, since neuroimaging data can be highly variable across individuals due to anatomical, functional, and cognitive differences (Dubois & Adolphs, 2016), counterbalancing was employed within participants. This approach ensured that each participant experienced different S-R mappings, thereby controlling for individual differences and increasing the sensitivity to detect true effects. Thus, participants completed the tasks twice, with a week between sessions, each session with different S-R mappings (i.e., blue–left, yellow–right, red–up, green–down, or blue–right, yellow–left, red–down, green–up). The order of S-R mappings varied across participants. To minimize potential confounding factors, each session was conducted on the same day of the week (e.g., every Monday), at the same time (e.g., at 2 PM), and under the same environmental conditions.

### Procedure

After providing informed consent, participants performed the tasks in a dimly-lit sound-proof chamber. The participant’s body midline and the numeric keypad were aligned with the center of the monitor. Participants were instructed to press one of the directional response keys on the numeric keypad depending on the color of the target stimulus as quickly and accurately as possible. The instructions also informed of the four S-R mappings. On each trial, when the fixation cross was presented, participants were instructed to press the home key with their right index finger and to keep pressing the key until they decided their response to the target after it was presented. 500 ms after the home key was pressed, a target stimulus was presented for 250 ms or until the home key was released. After the target display, a blank display was presented for 1,750 ms. Participants were instructed to press one of the four direction keys depending on the color of the target stimulus. All responses were executed using the right index finger. If the release of the home key occurred before target presentation, a visual feedback message was displayed, saying “Press the home key”. Each time a participant responded incorrectly or failed to respond within 2,000 ms after the target onset, a 150 ms beep sound was played as feedback.

Total reaction time (TT) was measured separately by initiation time (IT) and movement time (MT; Figure 2B). IT is defined as the time from the target onset to the moment when the home key is released. MT was defined as the time from the moment when the home key is released to the moment when a directional key is pressed. Evidence of IT CSE may reflect conflict resolution occurred at relatively early information processing stage (e.g., stimulus identification to response selection stage), whereas evidence of MT CSE may reflect conflict resolution at relatively late information processing stage (e.g., movement execution stage). It is important to note that some perspectives suggest response selection and execution are not entirely distinct processes, meaning responses can be initiated before response selection is fully completed (Buetti & Kerzel, 2008; Buetti & Kerzel, 2009; Calderon et al., 2018; Erlhagen & Schöner, 2002; Hommel, 2009; N. Lee & Cho, 2024; Resulaj et al., 2009). However, to track the temporal dynamics of conflict resolution in the behavioral data, we chose to separate TT into IT and MT, which allows for a more direct comparison with decoding results.

After a practice block of 34 trials, participants completed eight blocks of 82 trials, with 1-min breaks between blocks. The four sequential trial types of congruent trials followed by a congruent trial (cC), congruent trials followed by an incongruent trial (iC), incongruent trial followed by a congruent trial (cI), and incongruent trials followed by an incongruent trial) were equally frequently presented in the horizontal and vertical Simon tasks respectively. The congruency for the first and second trials of each block was randomly determined.

### EEG recording and preprocessing

EEG signals were recorded using SynAmps RT NeuroScan 64-channel EEG system (NeuroScan Compumedics, Charlotte, NC, USA) with a pass-band of 0.01 – 200 Hz. The ground electrode was positioned between the FPz and Fz electrodes. Signals from two additional electrodes attached to the left and right earlobes were used as the reference electrodes. Eye movements were measured with vertical (VEOG) and horizontal electrooculogram (HEOG). VEOG electrodes were positioned above and below the left eye, while HEOG electrodes were positioned 1 cm lateral to the outer canthi of each eye. Electrocardiogram (ECG) signals were recorded to exclude effects of cardiac signals (Park et al., 2014) with two electrodes placed approximately 2 cm below the left and right collarbones. Channel impedances were maintained below 15 kΩ.

The preprocessing of the signals was conducted using the EEGLAB Toolbox (Delorme & Makeig, 2004) and the ERPLAB Toolbox (Lopez-Caldeeron & Luck 2014). Scalp EEG signals were offline referenced to the average of the left and right earlobes. All signals were band-pass filtered (non-causal Butterworth impulse response function, half amplitude cutoffs at 0.1 and 80 Hz, 12 dB/oct roll-off) and resampled at 250 Hz. Any EEG signals containing significant muscle artifacts or extreme voltage offsets were automatically and manually identified and removed. Independent component analysis (ICA) was subsequently applied on the scalp EEG for each participant to identify and remove components associated with blinks (Jung et al., 2000) and eye movements (Drisdelle et al., 2017). Following ICA correction, the EEG data were segmented for each trial from −500 to +1,200 ms relative to the onset of the stimulus (i.e., baseline: [−500, 0]; stimulus presentation: [0, 250]; Blank: [250, 1200]). Standard artifact rejection procedures (Luck, 2014) were then implemented to eliminate epochs containing miscellaneous artifactual voltage deflection. The average percentage of rejected trials was 7.52%. After removing artifacts, the counterbalanced data for each participant were concatenated.

### Decoding Analysis

The preprocessed dataset was organized based on the previous congruency. Subsequently, the data were further divided according to the current color or location attributes. Thus, dataset was categorized into four groups: color information on trials following a congruent trial collapsed across the locations, location information following a congruent trial collapsed across the colors, color information following an incongruent trial collapsed across the locations, and location information following an incongruent trial collapsed across the colors. Within these categories, color information was labeled according to the color of the square of the current trial (i.e., blue-1, yellow-2, red-3, green-4). Similarly, location information were labeled based on the location of the square of the current trial (i.e., left-1, right-2, up-3, down-4).

In the decoding analysis, data were divided into two conditions (trials after congruent and trials after incongruent). Current congruency was intentionally collapsed within the dataset to avoid redundancy between the color and location datasets. If the data were divided into four conditions (cC, cI, iC, and iI), trials that are current congruent (e.g., a blue square on the left side either in cC and iC) would result in identical markings for both color and location datasets (e.g., both being labeled as “1”). This overlap would reduce the independence needed for effective analysis. To maintain a clear distinction between the color and location datasets, the data were divided based on previous congruency, while collapsing current congruency. This approach prevents the issue of identical labeling that would occur when mixing congruent and incongruent current trials. Similar methods of collapsing current congruency to focus on the effects of previous experiences have been applied in previous neuroimaging research (Pastötter et al., 2013; Fan et al., 2007).

For the decoding analysis, the Amsterdam Decoding and Modeling toolbox (Fahrenfort et al., 2018) was used on the preprocessed EEG data. Before classification, the EEG data were resampled to 50 Hz to reduce computation time. To ensure that the training of the classifier was not biased across classes, an undersampling method was applied. This involved randomly selecting trials from the conditions with more trials to balance them with the conditions having fewer trials, resulting in an equal count of subsequent congruent and subsequent incongruent trials within each class. All channels were used for decoding analysis. Training and testing were performed on the same dataset for each condition separately using a 5-fold cross-validation procedure. The dataset was divided into five equal-size groups of trials; four out of these groups were used for training and the remaining fifth was reserved for testing. Linear discriminant analysis (LDA) classifiers were adopted for each category of information, with each classifier trained to differentiate between one attribute versus the others. For example, in color decoding, models classified one color (e.g., red) versus the other colors, whereas in location decoding, the model classified one location (e.g., left) versus the other locations. The trained models were then employed to predict the color or location of each test data point reserved for testing. This process was repeated five times until all data were tested. Classifier performance was then averaged across folds.

It is important to note that the LDA classifier is unlikely to misinterpret color signals as location signals because location attributes are mixed within each color attribute, and vice versa. Additionally, the LDA classifier is unlikely to decode color or location attributes based solely on horizontal or vertical information, as both dimensions are integrated within each attribute. While some stimulus attributes are associated with specific dimensions (e.g., horizontal with blue and yellow, vertical with red and green), it is improbable that the classifier would differentiate attributes based purely on this. Even if dimensional information were used, the classifier would still need to detect distinct features within each dimension to accurately differentiate stimulus attributes. Finally, it is unlikely that the evidence for color decoding is confounded with response decoding, given that the mapping of colors to responses was strictly counterbalanced.

The area under the curve (AUC) per time point, derived from the signal detection theory, was used as a measure of classification accuracy. t-tests were conducted across participants against a 50% chance level. Although the model classifies one out of four attributes, the ADAM toolbox computes AUC for multi-class problems by averaging the AUC across all pairwise comparisons between classes (Fahrenfort et al., 2018). As a result, the chance AUC performance is consistently 50%, regardless of the number of classes being analyzed. Differences in classification accuracy between previous congruent and previous incongruent conditions were then calculated for both color and location accuracy. Cluster-based permutation tests (*p <* .05, 1,000 iterations) were then used to perform multiple-comparison correction for these t tests over time (Cohen, 2014; Maris & Oostenveld, 2007; Nichols & Holmes, 2002). The null distribution of the cluster size under random permutation was determined based on the observed cluster size, and p values for the clusters were calculated using this comparison. Since the neural representation of color or location starts from target onset and ends when the response is made, the analysis period for the decoding AUC was set from 0 to 600 ms. This duration was determined based on the average total reaction time, which was 508 ms (Table 1). This AUC reflects the accuracy of machine learning in discriminating stimulus attributes, which, in other words, indicates the strength of the stimulus representation in the brain. Finally, this decoding accuracy was calculated in different frequency bands: in theta (4-8 Hz), alpha (8-12 Hz), low beta (12-20 Hz), and high beta (20-30 Hz).

**Table 1.** Mean of total reaction time (TT), initiation time (IT), movement time (MT), and percentage errors (PE).

| N-1 cong | N-0 cong | TT | IT | MT | PE |
| --- | --- | --- | --- | --- | --- |
| Congruent | Congruent | 486 | 368 | 117 | 2.1 |
|  | Incongruent | 536 | 396 | 130 | 4.9 |
| Incongruent | Congruent | 495 | 376 | 118 | 2.2 |
|  | Incongruent | 525 | 398 | 126 | 3.3 |
| Average |  | 508 | 384 | 123 | 3.3 |

### Time-Frequency analysis

Time-frequency analysis was conducted to determine whether consistent results could be observed in comparison to the decoding outcomes. The data were decomposed into frequency bands using complex Morlet wavelet convolution, following the method described by Cohen (2014). The wavelets’ frequencies ranged from 2 to 40 Hz, comprising 60 linearly spaced wavelets. Complex Morlet wavelets were created by multiplying a Gaussian 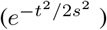,where *s* is the width of the Gaussian) with sine waves (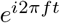, where *i* is the complex operator, *f* is frequency, and *t* is time). The Gaussian width was set as *s* = *δ/*(2*πf*), where *δ* represents the number of cycles of each wavelet, logarithmically spaced between 4 and 40 to have a good tradeoff between temporal and frequency precision. Frequency-domain convolution was applied, that is, applying the fast Fourier transform to both the EEG data and the Morlet wavelets, multiplying them, and converting the result back to the time domain using the inverse fast Fourier transform. The squared magnitude of these complex signals was taken at each time point and each frequency to acquire power, that is, [real(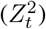) + imag(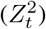)]. Power were then decibel normalized [dB Power_*tf*_ = 10\**log*10(Power_*tf*_ / Baseline Power_*f*_)], where baseline is the frequency-specific average of power values over −300 to −100 pre-stimulus time period.

Normalized power data were averaged based on the conditions; congruent trials following congruent trials (cC), incongruent trials following congruent trials (cI), congruent trials following incongruent trials (iC), and incongruent trials following incongruent trials (iI). The calculation for condition subtraction was based on the formula: CSE = (cI - cC) - (iC - iI). Statistics were performed by one sample t-tests against zero, and multiple comparisons were corrected through cluster-based permutation testing. In the permutation test, condition labels were randomly shuffled for each data point, and t-values were re-computed. The sum of t-values within the largest cluster was recorded into a distribution of summed cluster t-values. This process was repeated 1,000 times, generating a distribution of maximum cluster sizes under the null hypothesis. Any clusters in the true data that were greater than or less than 95% of the null distribution (i.e. *p <*.05) were considered statistically significant.

Time-frequency region of interest (ROI) analysis was further adopted to elucidate the difference between the conditions. The time-frequency ROI windows were selected based on the location of the cluster and decoding results, with the time window set at 400 to 500 ms. Power was then calculated separately for the combination of previous-trial congruency (congruent, incongruent) and current-trial congruency (congruent, incongruent). Two-way repeated measures ANOVAs were conducted on mean power with the above variables as within-subject variables.

The selection of electrodes was conducted independently of any potential variations across task conditions or frequency bands, ensuring an unbiased approach. First, we hypothesized that conflict resolution would most likely be observed at electrodes frequently cited in the literature for top-down conflict resolution, such as the frontocentral electrode FCz (Cohen & Donner, 2013; Duprez et al., 2020; Gulbinaite et al., 2014; Hoppe et al., 2017; Spape et al., 2011; van Driel et al., 2012; Wang et al., 2018) and the central electrode Cz (Chen & Melara, 2009; Clayson & Larson, 2011; Frühholz et al., 2011; Larson & Clayson, 2011; Rey-Mermet et al., 2019; Li et al., 2015; Tang et al., 2013). However, previous studies have also reported conflict resolution activity in other regions, including frontal, central-parietal, and posterior areas (Kalamala et al., 2020; Eder et al., 2012; Chen & Melara, 2009). To ensure a comprehensive evaluation, we analyzed condition-difference time-frequency topographical maps, ROI analysis, and conducted cluster-corrected permutation tests across all channels. This evaluation was performed in different frequency bands: theta (4-8 Hz), alpha (8-12 Hz), low beta (12-20 Hz), and high beta (20-30 Hz). The results revealed that Cz was the only electrode to exhibit a significant CSE and confirmed by cluster-corrected permutation testing. As a result, FCz was excluded from further analysis, and Cz was selected for subsequent time-frequency analyses. The lack of significant differences at FCz may be attributed to individual anatomical differences (Ignatiadis et al., 2022) or ethnic variations in cranial shape, such as brachycephaly (short-headed) and dolichocephaly (long-headed; Ball et al., 2010), which could cause slight variations in vertical electrode placement.

## Results

### Behavioral results

#### Total Time (TT)

The main effect of previous-trial congruency was significant, *F* (1, 29) = 9.59, *p* = .004, *MSE* = 46, 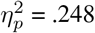, with the mean TT significantly greater after incongruent trials (*M* = 510 ms) than after congruent trials (*M* = 506 ms). A significant Simon effect was observed, as the main effect of current-trial congruency was significant,*F* (1, 29) = 153.14, *p <* .001, *MSE* = 242, 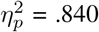. The mean TT was greater on incongruent trials (*M* = 525 ms) than congruent trials (*M* = 490 ms). There was a significant interaction between previous-trial congruency and current-trial congruency, *F* (1, 29) = 27.91, *p <* .001, *MSE* = 28, 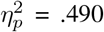, indicating a CSE (Figure 2C). The magnitude of the Simon effect was reduced after incongruent trials (30 ms), *F* (1, 29) = 124.91, *p <* .001, *MSE* = 108, 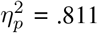, compared to after congruent trials (40 ms), *F* (1, 29) =149.72, *p <* .001, *MSE* = 163, 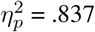.

#### Initiation Time (IT)

The main effect of previous-trial congruency was significant, *F* (1, 29) = 18.03, *p* = .002, *MSE* = 36, 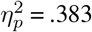, with the mean IT significantly greater after incongruent trials (*M* = 387 ms) than after congruent trials (*M* = 382 ms). The main effect of current-trial congruency was significant, *F* (1, 29) = 111.62, *p <* .001, *MSE* = 162, 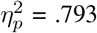. The mean IT was greater on incongruent trials (*M* = 397 ms) than congruent trials (*M* = 372 ms). The interaction between previous-trial congruency and current-trial congruency was significant, *F* (1, 29) = 8.87, *p* = .005, *MSE* = 25, 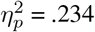 (Figure 2C). The magnitude of the Simon effect was reduced after incongruent trials (21 ms), *F* (1, 29) = 106.56, *p <* .001, *MSE* = 67, 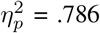, compared to after congruent trials (27 ms), *F* (1, 29) = 92.47, *p <* .001, *MSE* = 121, 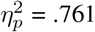. However, as the iI (*M* = 398 ms) was not faster than cI (*M* = 396 ms), this is not a typical pattern of the CSE (Figure 2).

#### MovementTime (MT)

The main effect of previous-trial congruency was marginally significant, *F* (1, 29) = 3.26, *p* = .081, *MSE* = 5.85, 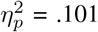. The mean MT was slightly lower after incongruent trials (*M* = 122 ms) than after congruent trials (*M* = 123 ms). The main effect of current-trial congruency was significant, *F* (1, 29) = 60.36, *p <* .001, *MSE* = 55, 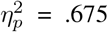. The mean MT was greater on incongruent trials (*M* = 128 ms) than congruent trials (*M* = 117 ms). The interaction between previous-trial congruency and current-trial congruency was significant, *F* (1, 29) = 19.8, *p <* .001, *MSE* = 8.8, 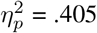, indicating a CSE (Figure 2C). The magnitude of the Simon effect was smaller after incongruent trials (8 ms), *F* (1, 29) = 40.82, *p <* .001, *MSE* = 24, 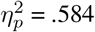, than after congruent trials (13 ms), *F* (1, 29) = 63.47, *p <* .001, *MSE* = 40, 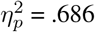.

#### Percentage Errors (PE)

The main effect of previous-trial congruency was not significant, *F* (1, 29) = 2.14, *p* = .15. The main effect of current-trial congruency was significant, *F* (1, 29) = 21.03, *p <* .001, *MSE* = 8.66, 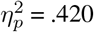. PE was higher on incongruent trials (4.66%) than congruent trials (2.20%). Interaction between previous-trial congruency and current-trial congruency was not significant, *F* (1, 29) = 1.9, *p* = .178 (Figure 2C).

### Decoding results

#### Theta (4-8 Hz)

The color decoding accuracy began to rise above chance level after the target onset, and gradually decreased after reaching its peak at approximately 200 ms but remained significant for the rest of the analysis period for both after congruent (cluster-corrected *p <*.05) and after incongruent trials (cluster-corrected *p <*.05; Figure 3A left).

Similarly, the location decoding accuracy began to rise above chance level after the target onset, and gradually decreased after reaching its peak at approximately 200 ms but remained significant for the rest of the analysis period for both after congruent (cluster-corrected *p <*.05) and after incongruent trials (cluster-corrected *p <*.05; Figure 3A right). However, there was no significant AUC difference between after congruent and incongruent trials in both color and location decoding (*p >* .05).

**Figure 3.**
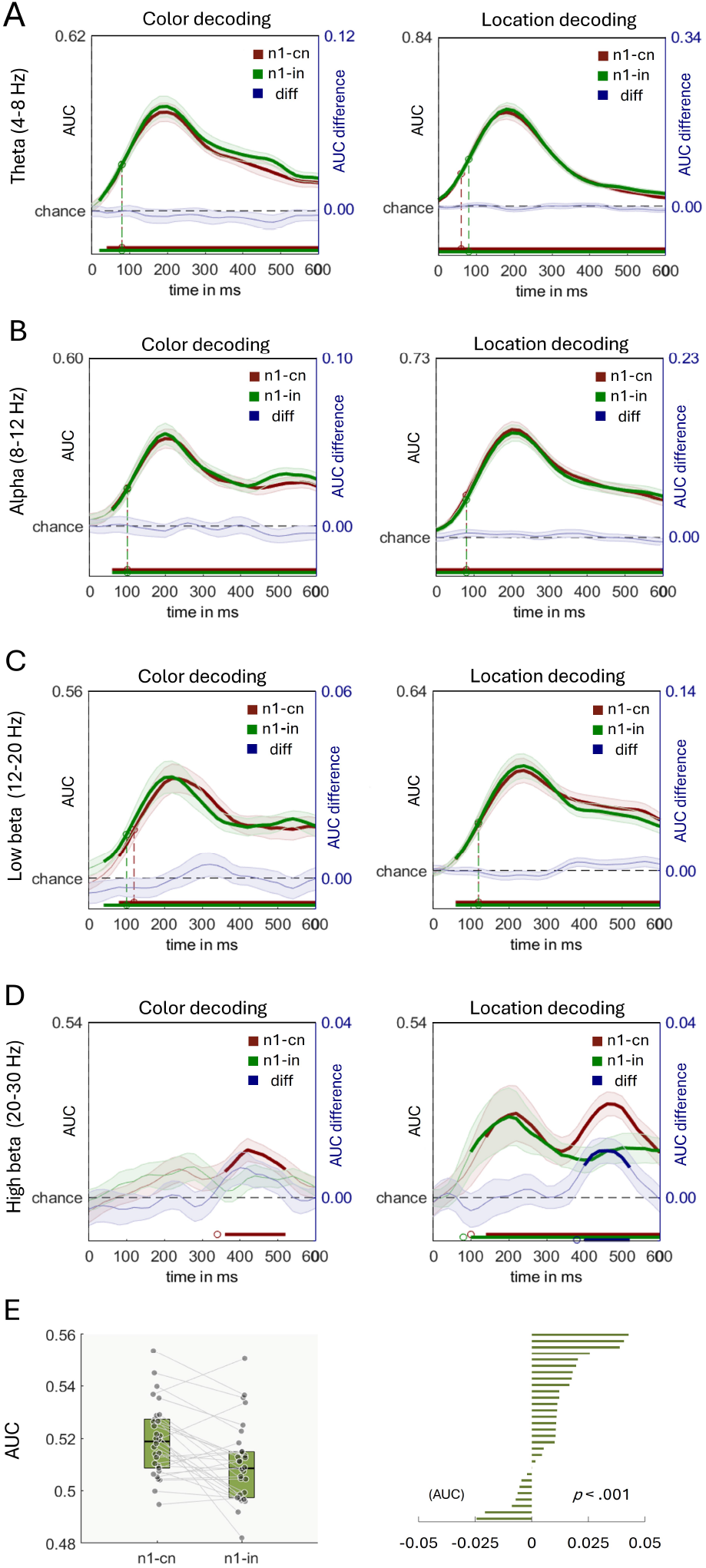
Decoding results. Color and location decoding AUCs were measured from target onset to response execution across different frequency bands: A. theta (4-8 Hz), B. alpha (8-12 Hz), C. low beta (12-20 Hz), and D. high beta (20-30 Hz). Thicker lines represent time windows where classification performance was statistically significant (*p <*.05, cluster-corrected). Here, “n1-cn” means previous congruent, “n1-in” means previous incongruent, “diff” means the accuracy differences between previous congruent and previous incongruent conditions. E. The left figure shows location decoding accuracy of previous congruent and previous incongruent trials in high beta frequency band (20-30 Hz) from 400 to 520 ms. The right figure shows individual congruency effect (i.e., previous congruent – previous incongruent) within the selected time and frequency window.

#### Alpha (8-12 Hz)

The color decoding accuracy began to rise above chance level after the target onset, and gradually decreased after reaching its peak at approximately 200 ms but remained significant for the rest of the analysis period for both after congruent (cluster-corrected *p <* .05) and after incongruent trials (cluster-corrected *p <* .05; Figure 3B left). Similarly, the location decoding accuracy began to rise above chance level after the target onset, and gradually decreased after reaching its peak at approximately 200 ms but remained significant for the rest of the analysis period for both after congruent (cluster-corrected *p <*.05) and after incongruent trials (cluster-corrected *p <* .05; Figure 3B right). However, there was no significant AUC difference between after congruent and incongruent trials in both color and location decoding (*p >* .05).

#### Low Beta (12-20 Hz)

The color decoding accuracy began to rise above chance level after the target onset, and gradually decreased after reaching its peak at approximately 250 ms but remained significant for the rest of the analysis period for both after congruent (cluster-corrected *p <* .05) and after incongruent trials (cluster-corrected *p <* .05; Figure 3C left). Similarly, the location decoding accuracy began to rise above chance level after the target onset, and gradually decreased after reaching its peak at approximately 200 ms but remained significant for the rest of the analysis period for both after congruent (cluster-corrected *p <*.05) and after incongruent trials (cluster-corrected *p <* .05; Figure 3C right). However, there was no significant AUC difference between after congruent and incongruent trials in both color and location decoding (*p >* .05).

#### High Beta (20-30 Hz)

There was significant above chance color decoding were observed in previous congruent trials (approximately from 350 ms to 505 ms, cluster-corrected *p <* .05; Figure 3D left). No other significant was observed in color decoding including difference between previous congruent and incongruent trials.

For location decoding, after congruent trials, the accuracy began to rise above chance level at 100 ms after the target onset and gradually decreased after reaching its peak at approximately 200 ms. The accuracy re-rise and reach its peak again at approximately 480 ms and remained significant for the rest of the analysis period (cluster-corrected *p <*.05; Figure 3D right). Whereas after incongruent trials, the location decoding accuracy began to rise above chance level at 100 ms after the target onset and gradually decreased after reaching its peak at approximately 200 ms but remained significant for the rest of the analysis period (cluster-corrected *p <* .05). Importantly, there was a significant difference between AUC after congruent and incongruent trials (from 400 ms to 520 ms, cluster-corrected *p* = .017), suggesting significantly lower decoding accuracy after incongruent trials compared to after congruent trials. To further confirm the result in univariate analysis, we extracted data from significant interval (i.e., from 400 ms to 520 ms), and averaged in each individual to conducted one-way ANOVA with previous-trial congruency (congruent, incongruent) as a within-subject variable. We found a significant difference in previous-trial congruency, *F* (1, 29) = 13.83, *p <* .001, *MSE <* .001, 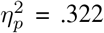 (Figure 3E), indicating a significant conflict modulation was observed in location decoding, particularly in high beta oscillations at 400 ms to 520 ms.

### Time-frequency results

#### Theta (4-8 Hz)

Electrodes FC1, FC2, F6, and FC6 showed statistically significant condition-related differences (*p <*.05; Figure 4A). To further explore the interaction pattern, we grouped FC1 and FC2 together, as well as F6 and FC6, based on the contour lines observed in the topographical map.

**Figure 4.**
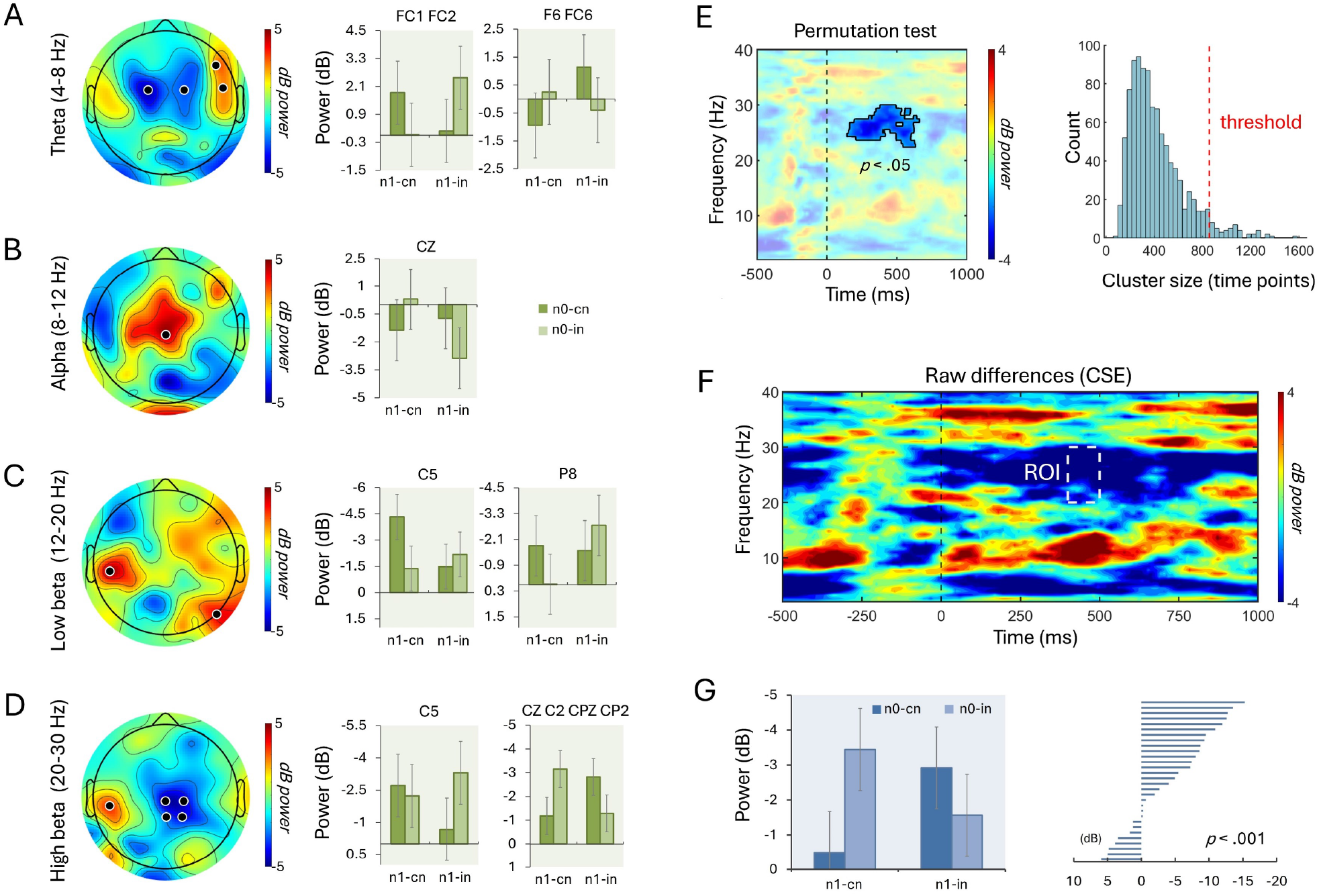
Time-frequency power results. Topographies show condition differences across different frequency bands: A. theta (4-8 Hz), B. alpha (8-12 Hz), C. low beta (12-20 Hz), and D. high beta (20-30 Hz). Black dots indicate electrodes with significant condition-related differences (*p <* .05). Electrodes were grouped based on topographical contours for further analysis. The bar graphs to the right display averaged data for four conditions (cC, cI, iC, and iI) in channels with significant effects from 400 to 500 ms. In the high beta band, averaged data from electrodes Cz, C2, CPz, and CP2 show a typical CSE pattern, indicating conflict resolution. Cz was ultimately selected for further analysis based on these results and previous studies. E. The cluster-based permutation test for Cz reveals a significant cluster on the z-map, with cluster size distribution shown alongside. F. Regions of interest (ROIs) are highlighted within the raw condition differences (i.e., CSE). G. presents averaged power for each condition (cC, cI, iC, and iI) from the ROI (left) and individual CSE results (right).

For the FC1 and FC2 group, the main effect of current-trial congruency was not significant, *F* (1, 29) *<* 1. However, a significant interaction between previous-trial congruency and current-trial congruency was observed, *F* (1, 29) = 9.55, *p* = .004, *MSE* = 13.12, 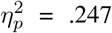. Similarly, for the F6 and FC6 group, the main effect of current-trial congruency was not significant, *F* (1, 29) *<* 1, but a significant interaction between previous-trial congruency and current-trial congruency was found, *F* (1, 29) = 5.73, *p* = .023, *MSE* = 9.81, 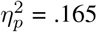. As depicted in Figure 4A, both interactions did not follow the typical pattern of the CSE (Figure 1). Furthermore, no significant clusters were found in any of the electrodes.

#### Alpha (8-12 Hz)

Electrode Cz exhibited statistically significant condition-related differences (*p <* .05; Figure 4B). The main effect of current-trial congruency was not significant, *F* (1, 29) *<* 1. However, a significant interaction between previous-trial congruency and current-trial congruency was observed, *F* (1, 29) = 5.68, *p* = .023, *MSE* = 19.16, 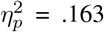. As shown in Figure 4B, this interaction did not follow the typical pattern of the CSE (Figure 1). Furthermore, no significant clusters were found in the alpha frequency domain.

#### Low Beta (12-20 Hz)

Electrodes C5 and P8 exhibited statistically significant condition-related differences (*p <* .05; Figure 4C). Based on the contour lines observed in the topographical map, we explored the interaction patterns of C5 and P8 separately. For C5, the main effect of current-trial congruency was not significant, *F* (1, 29) = 2.05, *p* = .162. However, a significant interaction between previous-trial congruency and current-trial congruency was found, *F* (1, 29) = 8.46, *p* = .006, *MSE* = 11.87, 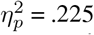. Additionally, a significant cluster was observed across conditions (cluster-corrected, *p <* .05), starting approximately 220 ms after stimulus presentation and continuing until 720 ms. Although a significant cluster was identified at C5, the interaction observed did not follow the typical pattern of the CSE (Figure 1). For P8, the main effect of current-trial congruency was not significant, *F* (1, 29) *<* 1. A significant interaction between previous-trial congruency and current-trial congruency was found, *F* (1, 29) = 4.7, *p* = .038, *MSE* = 14.02, 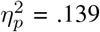. However, no clusters were observed at P8, and the interaction at this electrode also did not follow the typical CSE pattern (Figure 1).

#### High Beta (20-30 Hz)

Electrodes C5, Cz, C2, CPz, and CP2 exhibited statistically significant condition-related differences (*p <* .05; Figure 4D). Based on the contour lines observed in the topographical map, we analyzed the interaction patterns of C5 separately from the group of Cz, C2, CPz, and CP2. For C5, the main effect of current-trial congruency was not significant, *F* (1, 29) = 2.88, *p* = .100. However, a significant interaction between previous-trial congruency and current-trial congruency was found, *F* (1, 29) = 4.8, *p* = .036, *MSE* = 15.28, 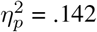. Despite this, no significant clusters were observed at C5, and the interaction did not follow the typical pattern of the CSE (Figure 1). For the group of Cz, C2, CPz, and CP2, the main effect of current-trial congruency was also not significant, *F* (1, 29) *<* 1. However, a significant interaction between previous-trial congruency and current-trial congruency was observed, *F* (1, 29) = 21.38, *p <* .001, *MSE* = 4.32, 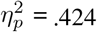. Importantly, the interaction pattern closely followed the typical CSE pattern (Figure 4D, Figure 1). Additionally, a significant cluster was observed across conditions (cluster-corrected, *p <* .05), beginning approximately 170 ms after stimulus presentation and continuing until 580 ms. Given that Cz is the electrode most commonly associated with top-down cognitive control in previous literature, we selected Cz for further time-frequency analysis.

#### Time-frequency analysis at CZ

In the cluster-based analysis, a significant reduction of power was observed in the high beta band (20-30 Hz) across conditions (cluster-corrected, *p <* .05; Figure 4E). This reduction started approximately 150 ms after stimulus presentation and continued until 640 ms. No other clusters were found in other time and frequency domains. For the ROI analysis, time-frequency windows were selected from 400 to 500 ms at 20-30 Hz. The main effect of previous-trial congruency was not significant, *F* (1, 29) *<* 1. The main effect of current-trial congruency was not significant, *F* (1, 29) = 1.17, *p* = .288. However, a significant interaction was obtained between previous-trial congruency and current-trial congruency, *F* (1, 29) = 14.06, *p <* .001, *MSE* = 9.85, 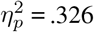, indicating a significant CSE was observed in high beta oscillations (Figure 4F,G). These results demonstrate that the decoding findings were validated through global time-frequency analysis.

## Discussion

In this study, we investigated how cognitive control resolves Simon conflicts and selectively processes information following incongruent trials. We decoded human scalp EEG data during Simon tasks to determine if brain representations of goal and distractor information change based on previous-trial congruency. In behavior analysis, we found a significant CSE in reaction times, indicating that top-down cognitive control was heightened after incongruent trials. Additionally, the CSE was observed in MT, suggesting that conflict resolution occurred during the movement execution stage. This stage encompasses the period from the release of the home key (averaging approximately 384 ms; Table 1) to the moment the actual response was made (averaging approximately 508 ms; Table 1). Decoding analysis demonstrated that the decoding accuracy of task-irrelevant location significantly decreased after incongruent trials compared to after congruent trials, particularly in the high beta frequency band (20-30 Hz) from approximately 400 ms to 520 ms. Conversely, no such difference was observed in the decoding of task-relevant color information.

Time-frequency analysis revealed a significant CSE in the high beta frequency band (20-30 Hz), further confirming that top-down cognitive control for Simon conflict resolution is associated with beta oscillations. These findings strongly support the idea that conflict resolution in Simon tasks is closely linked to the inhibition of task-irrelevant information processing and occurs during the movement execution stage.

The CSE is a hypothesized marker of top-down cognitive control which has been served as an important research tool for investigating the operation of the cognitive control. By testing whether the CSE transfers across different task conditions (e.g., conflict type, task sets, distractor type), researchers specify the scope of cognitive control and infer the mechanism involved in cognitive control processes (Braem et al., 2014). Two different types of mechanisms have been proposed (a) biasing stimulus processing (b) biasing response processing (Egner et al., 2007; Soutschek et al., 2013). Egner et al. (2007) suggested that these two types of conflict resolution processes depend on the source of conflict. In their combined Stroop-Simon color naming task, participants were instructed to respond to the ink color of a word presented to the left or right of fixation, while ignoring both the word’s meaning (Stroop conflict) and its location (Simon conflict). Notably, significant CSEs were observed within the same conflict type but not across different conflicts. The author suggested that the absence of the CSE between Stroop and Simon conflicts is due to independent control processes resolving each type of conflict. Specifically, Stroop conflict is resolved through early stimulus-biasing, enhancing the processing of task-relevant information, whereas Simon conflict is resolved through late response-biasing, involving the suppression of the task-irrelevant information processing. This distinction was further supported by fMRI findings, which showed that the resolution of Stroop conflict was associated with the recruitment of the superior parietal cortex, whereas the resolution of Simon conflict was associated with the recruitment of the ventral premotor cortex.

However, inconsistent with the source of conflict account, Akçay and Hazeltine (2008) found that the CSE was transferred across tasks only when the stimulus and response alternatives for two Simon tasks displayed on the same hemispace, but not when one Simon task was presented in the left and the other presented in the right hemispace.

The authors suggested that the degree to which participants perceive the Simon tasks as a single or different tasks determines whether the conflict is resolved by the same or different control processes. It has been suggested that this task representation or task-sets formed based on salient perceptual features, such as task-relevant features (Braem et al., 2014) or the sensory modality of task stimuli (Yang et al., 2017; Hazeltine et al., 2011), and the predictability of task (Grant et al., 2020). Therefore, if two tasks are separated in terms of a salient feature, participants use this salient feature to divide a complex task into two or more simpler tasks, or different task-sets to aid task performance (Grant et al., 2023). This emerging view of the CSE is known as the episodic retrieval account (Dignath et al, 2019; Grant et al., 2020; Grant et al., 2023; Weissman et al., 2016). The account suggests that participants form episodic memories of the task set, which includes information about stimuli, responses, context-defining features, and the relationships governed by task rules. When any of these contextual features are repeated in subsequent trials, they trigger the retrieval of the previous trial’s memory, leading control processes to adopt the same settings. This results in a reduced congruency effect following incongruent trials compared to congruent trials.

However, Y. Lee and Cho (2023) found that the CSE transferred across Simon tasks that were highly distinguished by sensory modality, task predictability, and task-relevant dimension, all of which are essential for differentiating a task set. This indicates that differences in these salient features do not determine the transfer of the CSE between Simon tasks. Since the only components shared between the tasks were the task-irrelevant dimension and response mode, which are critical for response suppression (Kim et al., 2015; J. Lee & Cho, 2013; Lim & Cho, 2021b; Stürmer et al., 2002), the authors concluded that conflict resolution in Simon tasks primarily occurs at the automatic route between the task-irrelevant dimension and response mode.

Consistent with Y. Lee and Cho (2023), our decoding results revealed that decoding accuracy for task-irrelevant information changed as a function of previous congruency, whereas no such modulation was observed for task-relevant information, indicating that conflict resolution in the Simon task primarily occurs at the task-irrelevant dimension. More importantly, decoding accuracy for task-irrelevant information was significantly reduced after incongruent trials compared to after congruent trials. This suggests that conflict resolution operated by suppressing the task-irrelevant information processing after experiencing conflict. Thus, building on the previous study that demonstrated the scope of conflict resolution in the Simon task (Y. Lee & Cho, 2023), our study is the first to provides direct evidence of how Simon conflicts are resolved through inhibition, which offers valuable insights into the mechanisms underlying top-down cognitive control.

Neural oscillations, often referred to as brain rhythms, are periodic patterns of electrical activity in the brain that arise from the synchronized activity of neurons (Fries, 2005). These oscillations are categorized into different frequency bands which are linked to distinct cognitive and behavioral functions (Buzsáki, 2006). Among these brain rhythms, oscillations in the beta frequency range (13-30 Hz) are particularly associated with sensorimotor processing (Barker, 2007; Engel & Fries, 2010; Pavlidou et al., 2014; Pfurtscheller et al., 1996; Schmidt et al., 2019; Spitzer & Haegens, 2017). During movement preparation and execution, beta power decreases, while an increase in beta power in sensorimotor areas reflects active suppression of the motor system. For example, modulations of sensorimotor beta power have been observed during the observation of correct versus incorrect movements, with decreased beta power for incorrect movements, suggesting that sensorimotor beta oscillations might be involved in evaluating observed movement (Koelewijn et al., 2008). Moreover, increased beta power at frontal scalp EEG electrodes is also associated with movement stopping (e.g., Swann et al., 2009; Swann et al., 2012). Using intracranial EEG, studies have reported increases in beta band activity in the right inferior frontal gyrus during successful stop trials compared to unsuccessful trials. These findings suggest that beta power may serve as a critical indicator of motor suppression.

Beyond its established role as a sensorimotor rhythm, beta oscillations have been implicated in a wide range of cognitive functions, primarily associated with top-down controlled processing (Buschman & Miller, 2007; Engel & Fries, 2010; Wang, 2010; Fries, 2015). According to the status-quo theory proposed by Engel and Fries (2010), beta oscillations are involved in maintaining a motor and/or cognitive state. Specifically, the theory proposed that the level of beta band activity remains constant when there is no change in the cognitive or perceptual set, increases when the current cognitive set has to be maintained, and decreases when the current setting is disrupted by a novel or unexpected event. Evidence supporting this theory comes from Buschman and Miller (2007), demonstrating a strong link between top-down processing and beta-band activity. In their study, monkeys were trained to find a target among distractors using either a pop-out mode (where the target is easily distinguishable from the distractors) or a serial search mode (where the target shares some features with distractors and is thus less distinguishable). They found that beta-band coherence between frontal and parietal regions was prevalent during serial searches, which required strong top-down processing. In contrast, gamma-band coupling was more significant during pop-out searches, which relied on bottom-up saliency of the target. These results suggest that beta oscillations are involved in endogenous top-down control, while gamma oscillations convey exogenous bottom-up signals. Consistent with this finding, numerous studies have also linked beta oscillations to top-down cognitive control (Axmacher et al., 2008; Deiber et al., 2007; Cunillera et al., 2012; Gladwin et al., 2006).

Our results further verify the strong association between beta oscillations and top-down cognitive control, specifically in terms of inhibition. Our time-frequency analysis revealed that beta power in the current trial significantly increased after experiencing incongruent trials compared to congruent trials, indicating that top-down control is heightened after experiencing conflict. Additionally, location decoding in high beta band was significantly decreased following incongruent trials compared to following congruent trials. Thus, by demonstrating top-down cognitive control in the CSE within the beta-frequency band in the context of a bottom-up confound-minimized paradigm, our findings provide clear evidence of this association. Importantly, since these effects were observed in the Simon task, our study raises the possibility that beta oscillations are specifically related to inhibition. Together, we interpret beta frequency in our task as the electrophysiological signature of the conflict resolution process, specifically reflecting response suppression activated by inhibitory control to prevent strong, unwanted habitual behaviors.

The absence of theta oscillation in the Simon task is noteworthy, as it contrasts with the extensive literature that emphasizes the importance of theta band activity in conflict resolution. We propose two potential explanations for this finding, primarily focusing on differences in experimental paradigms. Our confound-minimized design, in which no stimulus or response attributes were repeated across trials, may have emphasized inhibition more than other cognitive control functions, such as working memory and cognitive flexibility. Since inhibition is closely tied to sensorimotor processing, beta activity became more prominent in our study. Another possible explanation is the response method used in our task, which is more refined than traditional keypress methods. Previous research has shown that task responded with mouse tracking elicited beta oscillations (Palmer et al., 2019), which is similar to the present response method that divides responses into IT, MT, and TT. Thus, the use of an aimed movement method may have increased the demand for inhibition during task performance.

In conclusion, the present study demonstrates that the CSE in top-down cognitive control inherently involves inhibition, with its foundation resting on the suppression of distractor processing. This modulation is reflected in beta activity, which is recognized for its association with top-down cognitive control and motor inhibition. It is important to note that the present study does not claim to have completely ruled out other cognitive control processes that also contribute to sequential modulation in the Simon task. For instance, working memory is also an important component that influences Simon sequential modulation, depending on specific attributes of a given paradigm. Nevertheless, our findings highlight the important role of inhibition in the CSE, suggesting that the suppression of distractor representation is a key mechanism underlying Simon conflict resolution.

## Acknowledgement

This research was supported by the Korean Research Foundation Grant funded by the Korean Government (NRF-2020R1A2C2012033).

## Author Contributions

Yoon Seo Lee: Conceptualization; Data curation; Formal analysis; Investigation; Methodology; Project administration; Writing—Original draft; Writing—Review & editing. Gi-Yeul Bae. Conceptualization; Methodology; Writing—Review & editing. Yang Seok Cho: Conceptualization; Project administration; Writing—Review & editing.

## Notes

### Competing Interest Statement

The authors have declared no competing interest.

## References

Akçay, Ç., & Hazeltine, E. (2008). Conflict adaptation depends on task structure. Journal of Experimental Psychology: Human Perception and Performance, 34(4), 958–973. 10.1037/0096-1523.34.4.958

Arnau, S., Brümmer, T., Liegel, N., & Wascher, E. (2021). Inverse effects of time-on-task in task-related and task-unrelated theta activity. Psychophysiology, 58(6), Article e13805. 10.1111/psyp.13805

Axmacher, N., Schmitz, D. P., Wagner, T., Elger, C. E., & Fell, J. (2008). Interactions between medial temporal lobe, prefrontal cortex, and inferior temporal regions during visual working memory: a combined intracranial EEG and functional magnetic resonance imaging study. Journal of Neuroscience, 28(29), 7304–7312. 10.1523/JNEUROSCI.1778-08.2008

Bae, G. Y. (2021). The Time Course of Face Representations during Perception and Working Memory Maintenance. Cerebral Cortex, 2 (1), 1–12. 10.1093/texcom/tgaa093

Bae, G. Y., & Chen, K. W. (2024). EEG decoding reveals task-dependent recoding of sensory information in working memory. NeuroImage, 297, 120710. 10.1016/j.neuroimage.2024.120710

Bae, G. Y., & Luck, S. J. (2018). Dissociable decoding of spatial attention and working memory from EEG oscillations and sustained potentials. Journal of Neuroscience, 38(2), 409–422. 10.1523/JNEUROSCI.2860-17.2017

Bae, G. Y., & Luck, S. J. (2019). Reactivation of previous experiences in a working memory task. Psychological science, 30 (4), 587–595. 10.1177/0956797619830398

Ball, R., Shu, C., Xi, P., Rioux, M., Luximon, Y., & Molenbroek, J. (2010). A comparison between Chinese and Caucasian head shapes. Applied ergonomics, 41 (6), 832–839. 10.1016/j.apergo.2010.02.002

Baker, S. N. (2007). Oscillatory interactions between sensorimotor cortex and the periphery. Current opinion in neurobiology, 17(6), 649–655. 10.1016/j.conb.2008.01.007

Beatty, P. J., Buzzell, G. A., Roberts, D. M., Voloshyna, Y., & McDonald, C. G. (2021). Subthreshold error corrections predict adaptive post-error compensations. Psychophysiology, 58(6). 10.1111/psyp.13803

Botvinick, M. M., Braver, T. S., Barch, D. M., Carter, C. S., & Cohen, J. D. (2001). Conflict monitoring and cognitive control. Psychological Review, 108(3), 624–652. 10.1037/0033-295X.108.3.624

Braem, S., Abrahamse, E. L., Duthoo, W., & Notebaert, W. (2014). What determines the specificity of conflict adaptation? A review, critical analysis, and proposed synthesis.Frontiers in psychology, 5, 1134. 10.3389/fpsyg.2014.01134

Buetti, S., & Kerzel, D. (2008). Time course of the Simon effect in pointing movements for horizontal, vertical, and acoustic stimuli: Evidence for a common mechanism. Acta Psychologica, 129(3), 420–428. 10.1016/j.actpsy.2008.09.007

Buetti, S., & Kerzel, D. (2009). Conflicts during response selection affect response programming: reactions toward the source of stimulation. Journal of Experimental psychology. Human Perception and Performance, 35(3), 816–834. 10.1037/a0011092

Buschman, T. J., & Miller, E. K. (2007). Top-down versus bottom-up control of attention in the prefrontal and posterior parietal cortices. Science, 315(5820), 1860–1862. 10.1126/science.1138071

Buzsáki, G. (2006). Rhythms of the brain. Oxford University Press. 10.1093/acprof:oso/9780195301069.001.0001

Calderon, C. B., Gevers, W., & Verguts, T. (2018). The unfolding action model of initiation times, movement times, and movement paths. Psychological Review, 125(5), 785–805. 10.1037/rev0000110

Cao, Y., Cao, X., Yue, Z., & Wang, L. (2017). Temporal and spectral dynamics underlying cognitive control modulated by task-irrelevant stimulus-response learning. Cognitive, Affective, & Behavioral Neuroscience, 17, 158–173. 10.3758/s13415-016-0469-5

Chen, S., & Melara, R. D. (2009). Sequential effects in the Simon task: conflict adaptation or feature integration?. Brain Research, 1297, 89–100. 10.1016/j.brainres.2009.08.003

Chinn, L. K., Pauker, C. S., & Golob, E. J. (2018). Cognitive control and midline theta adjust across multiple timescales. Neuropsychologia, 111, 216–228. 10.1016/j.neuropsychologia.2018.01.031

Clayson, P. E., & Larson, M. J. (2011). Effects of repetition priming on electrophysiological and behavioral indices of conflict adaptation and cognitive control. Psychophysiology, 48(12), 1621–1630. 10.1111/j.1469-8986.2011.01265.x

Cohen, M. X. (2014). Analyzing neural time series data: Theory and practice. Cambridge, MA: MIT Press. 10.7551/mitpress/9609.001.0001

Cohen, M. X., & Donner, T. H. (2013). Midfrontal conflict-related theta-band power reflects neural oscillations that predict behavior. Journal of neurophysiology, 110(12), 2752–2763. 10.1152/jn.00479.2013

Cohen, M. X., & Ridderinkhof, K. R. (2013). EEG Source Reconstruction Reveals Frontal-Parietal Dynamics of Spatial Conflict Processing. PLOS ONE, 8(2), 1–14. 10.1371/journal.pone.0057293

Cunillera, T., Fuentemilla, L., Periañez, J., Marco-Pallarès, J., Krämer, U. M., Càmara, E., Münte, T. F., & Rodríguez-Fornells, A. (2012). Brain oscillatory activity associated with task switching and feedback processing. Cognitive, Affective & Behavioral Neuroscience, 12(1), 16–33. 10.3758/s13415-011-0075-5

De Jong, R., Liang, C.-C., & Lauber, E. (1994). Conditional and unconditional automaticity: A dual-process model of effects of spatial stimulus-response correspondence. Journal of Experimental Psychology: Human Perception and Performance, 20(4), 731–750. 10.1037/0096-1523.20.4.731

de Vries, I. E., Marinato, G., & Baldauf, D. (2021). Decoding object-based auditory attention from source-reconstructed MEG alpha oscillations. Journal of Neuroscience, 41(41), 8603–8617. 10.1523/JNEUROSCI.0583-21.2021

de Vries, I. E., Savran, E., van Driel, J., & Olivers, C. N. (2019). Oscillatory mechanisms of preparing for visual distraction. Journal of cognitive neuroscience, 31(12), 1873–1894. 10.1162/jocna01460

Deiber, M. P., Missonnier, P., Bertrand, O., Gold, G., Fazio-Costa, L., Ibanez, V., & Giannakopoulos, P. (2007). Distinction between perceptual and attentional processing in working memory tasks: a study of phase-locked and induced oscillatory brain dynamics. Journal of cognitive neuroscience, 19(1), 158–172. 10.1162/jocn.2007.19.1.158

Delorme, A., & Makeig, S. (2004). EEGLAB: an open source toolbox for analysis of single-trial EEG dynamics including independent component analysis. Journal of neuroscience methods, 134(1), 9–21. 10.1016/j.jneumeth.2003.10.009

Diamond, A. (2013). Executive functions. Annual review of psychology, 64(1), 135–168. 10.1146/annurev-psych-113011-143750

Dignath, D., Johannsen, L., Hommel, B., & Kiesel, A. (2019). Reconciling cognitive-control and episodic-retrieval accounts of sequential conflict modulation: Binding of control-states into event-files. Journal of Experimental Psychology: Human Perception and Performance, 45(9), 1265–1270. 10.1037/xhp0000673

Drisdelle, B. L., Aubin, S., & Jolicoeur, P. (2017). Dealing with ocular artifacts on lateralized ERPs in studies of visual-spatial attention and memory: ICA correction versus epoch rejection. Psychophysiology, 54(1), 83–99. 10.1111/psyp.12675

Dubois, J., & Adolphs, R. (2016). Building a science of individual differences from fMRI. Trends in cognitive sciences, 20(6), 425–443. 10.1016/j.tics.2016.03.014

Duprez, J., Gulbinaite, R., & Cohen, M. X. (2019). Midfrontal theta phase coordinates behaviorally relevant brain computations during cognitive control. Neuroimage, 207, 116340–116340. 10.1016/j.neuroimage.2019.116340

Duprez, J., Houvenaghel, J. F., Dondaine, T., Péron, J., Haegelen, C., Drapier, S., … & Sauleau, P. (2019). Subthalamic nucleus local field potentials recordings reveal subtle effects of promised reward during conflict resolution in Parkinson’s disease. Neuroimage, 197, 232–242. 10.1016/j.neuroimage.2019.04.071

Eder, A. B., Leuthold, H., Rothermund, K., & Schweinberger, S. R. (2012). Automatic response activation in sequential affective priming: An ERP study. Social Cognitive and Affective Neuroscience, 7(4), 436–445. 10.1093/scan/nsr033

Egner, T., Delano, M., & Hirsch, J. (2007). Separate conflict-specific cognitive control mechanisms in the human brain. Neuroimage, 35(2), 940–948. 10.1016/j.neuroimage.2006.11.061

Egner, T., & Hirsch, J. (2005). Cognitive control mechanisms resolve conflict through cortical amplification of task-relevant information. Nature Neuroscience, 8(12), 1784–1790. 10.1038/nn1594

Engel, A. K., & Fries, P. (2010). Beta-band oscillations—signalling the status quo?. Current opinion in neurobiology, 20(2), 156–165. 10.1016/j.conb.2010.02.015

Erlhagen, W., & Schöner, G. (2002). Dynamic Field Theory of Movement Preparation. Psychological Review, 109(3), 545–572. 10.1037/0033-295x.109.3.545

Fahrenfort, J. J., van Driel, J., van Gaal, S., & Olivers, C. N. L. (2018). From ERPs to MVPA Using the Amsterdam Decoding and Modeling Toolbox (ADAM). Frontiers in Neuroscience, 12, 368–368. 10.3389/fnins.2018.00368

Fan, J., Byrne, J., Worden, M. S., Guise, K. G., McCandliss, B. D., Fossella, J., & Posner, M. I. (2007). The relation of brain oscillations to attentional networks. Journal of Neuroscience, 27(23), 6197–6206. 10.1523/JNEUROSCI.1833-07.2007

Faul, F., Erdfelder, E., Buchner, A., & Lang, A. G. (2009). Statistical power analyses using G* Power 3.1: Tests for correlation and regression analyses. Behavior research methods, 41(4), 1149–1160. 10.3758/brm.41.4.1149

Foster, J. J., & Awh, E. (2019). The role of alpha oscillations in spatial attention: limited evidence for a suppression account. Current opinion in psychology, 29, 34–40. 10.1016/j.copsyc.2018.11.001

Frackowiak, R. S. J., Friston, K. J., Frith, C. D., Dolan, R. J., Price, C. J., Zeki, S., Ashburner, J. T., & Penny, W. D. (2004). Human Brain Function (Second edition). Elsevier Academic Press. 10.1016/B978-0-12-264841-0.X5000-8

Fries, P. (2005). A mechanism for cognitive dynamics: Neuronal communication through neuronal coherence. Trends in Cognitive Sciences, 9(10), 474–480. 10.1016/j.tics.2005.08.011

Fries, P. (2015). Rhythms for cognition: communication through coherence. Neuron, 88(1), 220–235. 10.1016/j.neuron.2015.09.034

Frühholz, S., Godde, B., Finke, M., & Herrmann, M. (2011). Spatio-temporal brain dynamics in a combined stimulus–stimulus and stimulus–response conflict task. Neuroimage, 54(1), 622–634. 10.1016/j.neuroimage.2010.07.071

Gladwin, T. E., Lindsen, J. P., & de Jong, R. (2006). Pre-stimulus EEG effects related to response speed, task switching and upcoming response hand. Biological psychology, 72(1), 15–34. 10.1016/j.biopsycho.2005.05.005

Grant, L. D., Cookson, S. L., & Weissman, D. H. (2020). Task sets serve as boundaries for the congruency sequence effect. Journal of Experimental Psychology: Human Perception and Performance, 46(8), 798–812. 10.1037/xhp0000750

Grant, L. D., & Weissman, D. H. (2023). The binary structure of event files generalizes to abstract features: A nonhierarchical explanation of task set boundaries for the congruency sequence effect. Journal of Experimental Psychology: Learning, Memory, and Cognition, 49(7), 1033–1050. 10.1037/xlm0001148

Gratton, G., Coles, M. G. H., & Donchin, E. (1992). Optimizing the use of information: Strategic control of activation of responses. Journal of Experimental Psychology: General, 121(4), 480–506. 10.1037/0096-3445.121.4.480

Gulbinaite, R., van Rijn, H., & Cohen, M. X. (2014). Fronto-parietal network oscillations reveal relationship between working memory capacity and cognitive control. Frontiers in Human Neuroscience, 8, 761–761. 10.3389/fnhum.2014.00761

Hazeltine, E., Lightman, E., Schwarb, H., & Schumacher, E. H. (2011). The boundaries of sequential modulations: Evidence for set-level control. ncJournal of Experimental Psychology: Human Perception and Performae, 37(6), 1898–1914. 10.1037/a0024662

Hommel, B., Proctor, R. W., & Vu, K. P. L. (2004). A feature-integration account of sequential effects in the Simon task. Psychological research, 68, 1–17. 10.1007/s00426-003-0132-y

Hommel, B. (2009). Action control according to TEC (theory of event coding). Psychological Research, 73, 512–526. 10.1007/s00426-009-0234-2

Hoppe, K., Küper, K., & Wascher, E. (2017). Sequential Modulations in a Combined Horizontal and Vertical Simon Task: Is There ERP Evidence for Feature Integration Effects?. Frontiers in Psychology, 8, 1094–1094. 10.3389/fpsyg.2017.01094

Ignatiadis, K., Barumerli, R., Tóth, B., & Baumgartner, R. (2022). Effects of individualized brain anatomies and EEG electrode positions on inferred activity of the primary auditory cortex. Frontiers in Neuroinformatics, 16, 970372. 10.3389/fninf.2022.970372

Jung, T. P., Makeig, S., Westerfield, M., Townsend, J., Courchesne, E., & Sejnowski, T. J. (2000). Removal of eye activity artifacts from visual event-related potentials in normal and clinical subjects. Clinical neurophysiology, 111(10), 1745-1758. 10.1016/S1388-2457(00)00386-2

Kalamala, P., Ociepka, M., & Chuderski, A. (2020). ERP evidence for rapid within-trial adaptation of cognitive control during conflict resolution. Cortex, 131, 151–163. 10.1016/j.cortex.2020.07.012

Kerns, J. G., Cohen, J. D., MacDonald, A.W. III, Cho, R. Y., Stenger, V. A., & Carter, C. S. (2004). Anterior Cingulate Conflict Monitoring and Adjustments in Control. Science, 303(5660), 1023–1026. 10.1126/science.1089910

Kim, S., & Cho, Y. S. (2014). Congruency sequence effect without feature integration and contingency learning. Acta psychologica, 149, 60–68. 10.1016/j.actpsy.2014.03.004

Kim, S., Lee, S. H., & Cho, Y. S. (2015). Control processes through the suppression of the automatic response activation triggered by task-irrelevant information in the Simon-type tasks. Acta Psychologica, 162, 51–61. 10.1016/j.actpsy.2015.10.001

Koelewijn, T., van Schie, H. T., Bekkering, H., Oostenveld, R., & Jensen, O. (2008). Motor-cortical beta oscillations are modulated by correctness of observed action. Neuroimage, 40(2), 767–775.10.1016/j.neuroimage.2007.12.018

Koob, V., Mackenzie, I., Ulrich, R., Leuthold, H., & Janczyk, M. (2022). The role of task-relevant and task-irrelevant information in congruency sequence effects: Applying the diffusion model for conflict tasks. Cognitive Psychology, 140, 101528–101528. 10.1016/j.cogpsych.2022.101528

Kornblum, S., Hasbroucq, T., & Osman, A. (1990). Dimensional overlap: cognitive basis for stimulus-response compatibility–a model and taxonomy. Psychological review, 97(2), 253–270. 10.1037/0033-295x.97.2.253

Larson, M. J., & Clayson, P. E. (2011). The relationship between cognitive performance and electrophysiological indices of performance monitoring. Cognitive, Affective, Behavioral Neuroscience, 11, 159–171. 10.3758/s13415-010-0018-6

Lee, J., & Cho, Y. S. (2013). Congruency sequence effect in cross-task context: evidence for dimension-specific modulation. Acta Psychologica, 144(3), 617–627. 10.1016/j.actpsy.2013.09.013

Lee, N., & Cho, Y. S. (2024). Investigating the nature of spatial codes for different modes of Simon tasks: Evidence from congruency sequence effects and delta functions. Journal of Experimental Psychology: Human Perception and Performance, 50(8), 819–841.10.1037/xhp0001220

Lee, Y. S., & Cho, Y. S. (2023). The congruency sequence effect of the Simon task in a cross-modality context. Journal of Experimental Psychology: Human Perception and Performance, 49(9), 1221–1235. 10.1037/xhp0001145

Li, Q., Wang, K., Nan, W., Zheng, Y., Wu, H., Wang, H., & Liu, X. (2015). Electrophysiological dynamics reveal distinct processing of stimulus-stimulus and stimulus-response conflicts. Psychophysiology, 52(4), 562–571. 10.1111/psyp.12382

Li, Q., Yin, S., Wang, J., Zhang, M., Li, Z., Chen, X., & Chen, A. (2023). Not all errors are created equal: decoding the error-processing mechanisms using alpha oscillations. Cerebral Cortex, 33(13), 8110–8121. 10.1093/cercor/bhad102

Lim, C. E., & Cho, Y. S. (2018). Determining the scope of control underlying the congruency sequence effect: roles of stimulus-response mapping and response mode. Acta psychologica, 190, 267–276. 10.1016/j.actpsy.2018.08.012

Lim, C. E., & Cho, Y. S. (2021a). Cross-task congruency sequence effect without the contribution of multiple expectancy. Acta Psychologica, 214, Article 103268. 10.1016/j.actpsy.2021.103268

Lim, C. E., & Cho, Y. S. (2021b). Response mode modulates the congruency sequence effect in spatial conflict tasks: Evidence from aimed-movement responses. Psychological Research, 85(5), 2047–2068. 10.1007/s00426-020-01376-3

Lopez-Calderon, J., & Luck, S. J. (2014). ERPLAB: an open-source toolbox for the analysis of event-related potentials. Frontiers in Human Neuroscience, 8, 213–213. 10.3389/fnhum.2014.00213

Luck, S. J. (2014). An introduction to the event-related potential technique (Second edition). The MIT Press.

Maris, E., & Oostenveld, R. (2007). Nonparametric statistical testing of EEG-and MEG-data. Journal of neuroscience methods, 164(1), 177–190.10.1016/j.jneumeth.2007.03.024

Mayr, U., Awh, E., & Laurey, P. (2003). Conflict adaptation effects in the absence of executive control. Nature neuroscience, 6(5), 450–452. 10.1038/nn1051

Miyake, A., Friedman, N. P., Emerson, M. J., Witzki, A. H., Howerter, A., & Wager, T. D. (2000). The unity and diversity of executive functions and their contributions to complex “frontal lobe” tasks: A latent variable analysis. Cognitive psychology, 41(1), 49–100. 10.1006/cogp.1999.0734

Munakata, Y., Herd, S. A., Chatham, C. H., Depue, B. E., Banich, M. T., & O’Reilly, R. C. (2011). A unified framework for inhibitory control. Trends in cognitive sciences, 15(10), 453–459. 10.1016/j.tics.2011.07.011

Nichols, T. E., & Holmes, A. P. (2002). Nonparametric permutation tests for functional neuroimaging: a primer with examples. Human brain mapping, 15(1), 1–25. 10.1002/hbm.1058

Notebaert, W., & Verguts, T. (2008). Cognitive control acts locally. Cognition, 106(2), 1071–1080. 10.1016/j.cognition.2007.04.011

Notebaert, W., Gevers, W., Verbruggen, F., & Liefooghe, B. (2006). Top-down and bottom-up sequential modulations of congruency effects. Psychonomic bulletin & review, 13(1), 112–117. 10.3758/bf03193821

Nyhus, E., & Curran, T. (2010). Functional role of gamma and theta oscillations in episodic memory. Neuroscience & Biobehavioral Reviews, 34(7), 1023–1035. 10.1016/j.neubiorev.2009.12.014

Palmer, C. E., Auksztulewicz, R., Ondobaka, S., & Kilner, J. M. (2019). Sensorimotor beta power reflects the precision-weighting afforded to sensory prediction errors. Neuroimage, 200, 59–71. 10.1016/j.neuroimage.2019.06.034

Park, H. D., Correia, S., Ducorps, A., & Tallon-Baudry, C. (2014). Spontaneous fluctuations in neural responses to heartbeats predict visual detection. Nature neuroscience, 17(4), 612–618. 10.1038/nn.3671

Pashler, H. (1997). The psychology of attention. MIT Press. 10.7551/mitpress/5677.001.0001

Pavlidou, A., Schnitzler, A., & Lange, J. (2014). Beta oscillations and their functional role in movement perception. Translational Neuroscience, 5, 286–292. 10.2478/s13380-014-0236-4

Pastötter, B., Dreisbach, G., & Bäuml, K. H. T. (2013). Dynamic adjustments of cognitive control: oscillatory correlates of the conflict adaptation effect. Journal of Cognitive Neuroscience, 25(12), 2167–2178. 10.1162/jocna00474

Peylo, C., Hilla, Y., & Sauseng, P. (2021). Cause or consequence? Alpha oscillations in visuospatial attention. Trends in Neurosciences, 44(9), 705–713. 10.1016/j.tins.2021.05.004

Pfurtscheller, G., Stancak Jr, A., & Neuper, C. (1996). Post-movement beta synchronization. A correlate of an idling motor area?. Electroencephalography and clinical neurophysiology, 98(4), 281–293. 10.1016/0013-4694(95)00258-8

Resulaj, A., Kiani, R., Wolpert, D. M., & Shadlen, M. N. (2009). Changes of mind in decision-making. Nature, 461(7261), 263–266. 10.1038/nature08275

Rey-Mermet, A., Gade, M., & Steinhauser, M. (2019). Sequential conflict resolution under multiple concurrent conflicts: An ERP study. NeuroImage, 188, 411–418. 10.1016/j.neuroimage.2018.12.031

Riddle, J., Scimeca, J. M., Cellier, D., Dhanani, S., & D’Esposito, M. (2020). Causal evidence for a role of theta and alpha oscillations in the control of working memory. Current Biology, 30(9), 1748–1754. 10.1016/j.cub.2020.02.065

Schmidt, J. R., & De Houwer, J. (2011). Now you see it, now you don’t: Controlling for contingencies and stimulus repetitions eliminates the Gratton effect. Acta psychologica, 138(1), 176–186. 10.1016/j.actpsy.2011.06.002

Schmidt, R., Ruiz, M. H., Kilavik, B. E., Lundqvist, M., Starr, P. A., & Aron, A. R. (2019). Beta oscillations in working memory, executive control of movement and thought, and sensorimotor function. Journal of Neuroscience, 39(42), 8231–8238. 10.1523/JNEUROSCI.1163-19.2019

Schmidt, J. R., & Weissman, D. H. (2014). Congruency sequence effects without feature integration or contingency learning confounds. PLoS One, 9(7), 10.1371/journal.pone.0102337

Schneider, D., Herbst, S. K., Klatt, L. I., & Wöstmann, M. (2022). Target enhancement or distractor suppression? Functionally distinct alpha oscillations form the basis of attention. European Journal of Neuroscience, 55(11-12), 3256–3265. 10.1111/ejn.15309

Soutschek, A., Muller, H. J., & Schubert, T. (2013). Conflict-specific effects of accessory stimuli on cognitive control in the Stroop task and the Simon task. Experimental Psychology, 60(2), 140-147. 10.1027/1618-3169/a000181

Spapé, M. M., Band, G. P., & Hommel, B. (2011). Compatibility-sequence effects in the Simon task reflect episodic retrieval but not conflict adaptation: evidence from LRP and N2. Biological Psychology, 88(1), 116–123. 10.1016/j.biopsycho.2011.07.001

Spitzer, B., & Haegens, S. (2017). Beyond the status quo: a role for beta oscillations in endogenous content (re) activation. eneuro, 4(4). 10.1523/eneuro.0170-17.2017

Stürmer, B., Leuthold, H., Soetens, E., Schröter, H., & Sommer, W. (2002). Control over location-based response activation in the Simon task: Behavioral and electrophysiological evidence. Journal of Experimental Psychology: Human Perception and Performance, 28(6), 1345–1363. 10.1037/0096-1523.28.6.1345

Swann, N. C., Cai, W., Conner, C. R., Pieters, T. A., Claffey, M. P., George, J. S., … & Tandon, N. (2012). Roles for the pre-supplementary motor area and the right inferior frontal gyrus in stopping action: electrophysiological responses and functional and structural connectivity. Neuroimage, 59(3), 2860–2870. 10.1016/j.neuroimage.2011.09.049

Swann, N., Tandon, N., Canolty, R., Ellmore, T. M., McEvoy, L. K., Dreyer, S., … & Aron, A. R. (2009). Intracranial EEG reveals a time-and frequency-specific role for the right inferior frontal gyrus and primary motor cortex in stopping initiated responses. Journal of Neuroscience, 29(40), 12675–12685. 10.1523/JNEUROSCI.3359-09.2009

Tang, D., Hu, L., & Chen, A. (2013). The neural oscillations of conflict adaptation in the human frontal region. Biological Psychology, 93(3), 364–372. 10.1016/j.biopsycho.2013.03.004

Thut, G., Nietzel, A., Brandt, S. A., & Pascual-Leone, A. (2006). α-Band electroencephalographic activity over occipital cortex indexes visuospatial attention bias and predicts visual target detection. Journal of Neuroscience, 26(37), 9494–9502. 10.1523/JNEUROSCI.0875-06.2006

Töllner, T., Wang, Y., Makeig, S., Müller, H. J., Jung, T. P., & Gramann, K. (2017). Two independent frontal midline theta oscillations during conflict detection and adaptation in a Simon-type manual reaching task. Journal of Neuroscience, 37(9), 2504–2515. 10.1523/JNEUROSCI.1752-16.2017

Van Diepen, R. M., Foxe, J. J., & Mazaheri, A. (2019). The functional role of alpha-band activity in attentional processing: the current zeitgeist and future outlook. Current opinion in psychology, 29, 229–238. 10.1016/j.copsyc.2019.03.015

van Driel, J., Ort, E., Fahrenfort, J. J., & Olivers, C. N. (2019). Beta and theta oscillations differentially support free versus forced control over multiple-target search. Journal of Neuroscience, 39(9), 1733–1743. 10.1523/JNEUROSCI.2547-18.2018

van Driel, J., Ridderinkhof, K. R., & Cohen, M. X. (2012). Not all errors are alike: theta and alpha EEG dynamics relate to differences in error-processing dynamics. Journal of Neuroscience, 32(47), 16795–16806. 10.1523/JNEUROSCI.0802-12.2012

van Driel, J., Sligte, I. G., Linders, J., Elport, D., & Cohen, M. X. (2015). Frequency band-specific electrical brain stimulation modulates cognitive control processes. PLoS ONE, 10(9), Article e0138984. 10.1371/journal.pone.0138984

van Es, M. W., Gross, J., & Schoffelen, J. M. (2020). Investigating the effects of pre-stimulus cortical oscillatory activity on behavior. Neuroimage, 223, 117351. 10.1016/j.neuroimage.2020.117351

Wang, X. J. (2010). Neurophysiological and computational principles of cortical rhythms in cognition. Physiological reviews, 90(3), 1195–1268. 10.1152/physrev.00035.2008

Wang, X., Du, F., Hopfinger, J. B., & Zhang, K. (2018). Impaired conflict monitoring near the hands: Neurophysiological evidence. Biological psychology, 138, 41–47. 10.1016/j.biopsycho.2018.08.008

Wang, L., Chang, W., Krebs, R. M., Boehler, C. N., Theeuwes, J., & Zhou, X. (2019). Neural dynamics of reward-induced response activation and inhibition. Cerebral Cortex, 29(9), 3961–3976. 10.1093/cercor/bhy275

Weissman, D. H., Hawks, Z. W., & Egner, T. (2016). Different levels of learning interact to shape the congruency sequence effect. Journal of Experimental Psychology: Learning, Memory, and Cognition, 42(4), 566–583. 10.1037/xlm0000182

Wiesman, A. I., Koshy, S. M., Heinrichs-Graham, E., & Wilson, T. W. (2020). Beta and gamma oscillations index cognitive interference effects across a distributed motor network. Neuroimage, 213, 116747–116747. 10.1016/j.neuroimage.2020.116747

Yang, G., Nan, W., Zheng, Y., Wu, H., Li, Q., & Liu, X. (2017). Distinct cognitive control mechanisms as revealed by modality-specific conflict adaptation effects. Journal of Experimental Psychology: Human Perception and Performance, 43(4), 807–818. 10.1037/xhp0000351

